# DeepSimulator: a deep simulator for Nanopore sequencing

**DOI:** 10.1101/238683

**Authors:** Yu Li, Renmin Han, Chongwei Bi, Mo Li, Sheng Wang, Xin Gao

**Affiliations:** King Abdullah University of Science and Technology (KAUST), Computational Bioscience Research Center (CBRC), Computer, Electrical and Mathematical Sciences and Engineering (CEMSE) Division, Thuwal, 23955-6900, Saudi Arabia; King Abdullah University of Science and Technology (KAUST), Biological and Environmental Sciences and Engineering (BESE) Division, Thuwal, 23955-6900, Saudi Arabia

## Abstract

**Motivation:** Oxford Nanopore sequencing is a rapidly developed sequencing technology in recent years. To keep pace with the explosion of the downstream data analytical tools, a versatile Nanopore sequencing simulator is needed to complement the experimental data as well as to benchmark those newly developed tools. However, all the currently available simulators are based on simple statistics of the produced reads, which have difficulty in capturing the complex nature of the Nanopore sequencing procedure, the main task of which is the generation of raw electrical current signals.

**Results:** Here we propose a deep learning based simulator, DeepSimulator, to mimic the entire pipeline of Nanopore sequencing. Starting from a given reference genome or assembled contigs, we simulate the electrical current signals by a context-dependent deep learning model, followed by a base-calling procedure to yield simulated reads. This workflow mimics the sequencing procedure more naturally. The thorough experiments performed across four species show that the signals generated by our context-dependent model are more similar to the experimentally obtained signals than the ones generated by the official context-independent pore model. In terms of the simulated reads, we provide a parameter interface to users so that they can obtain the reads with different accuracies ranging from 83% to 97%. The reads generated by the default parameter have almost the same properties as the real data. Two case studies demonstrate the application of DeepSimulator to benefit the development of tools in *de novo* assembly and in low coverage SNP detection.

**Availability:** The software can be accessed freely at: https://github.com/lykaust15/deep_simulator.

## 1 INTRODUCTION

Next-generation sequencing (NGS) technologies allow researchers to sequence DNA and RNA in a high-throughput manner, which have facilitated numerous breakthroughs in genomics, transcriptomics, and epigenomics (Metzker, 2010; MacLean *et al*., 2009; Wu *et al*., 2017). The most popular NGS technologies on the market include Illumina, PacBio and Nanopore. Unlike the other sequencing technologies, Nanopore, whose core component is the pore chemistry that contains a voltage-biased membrane embedded with nanopores, would detect the electrical current signal changes when DNA or RNA molecules are forced to pass through the pore by voltage. Inputting the detected signals to a basecaller specifically designed for Nanopore, one can obtain the nucleotide sequence reads. Benefited from the underlying design, Nanopore sequencing owns the advantages of long-reads (Byrne *et al*., 2017), point-of-care (Lu *et al*., 2016), and PCR-free (Simpson *et al*., 2017), which enable *de novo* genome or transcriptome assembling with repetitive regions, field real-time analysis, and direct epigenetic detection, respectively.

Along with the rapid development in Nanopore sequencing, the downstream data analytical methods and tools have also been rapidly emerging. For example, Graphmap (Sović *et al*., 2016), Minimap2 (Li, 2017) and MashMap2 (Jain *et al*., 2017a) were particularly designed to map the Nanopore data to the genome. Canu (Koren *et al*., 2017) and Racon (Vaser *et al*., 2017) were created to assemble long and noisy reads produced by Nanopore. It is foreseeable that an even larger number of methods and tools would be developed in the near future. Therefore, it is quite important to benchmark those new methods using either empirical data (i.e., experimentally obtained) or simulated data (Escalona *et al*., 2016). Although it is essential that one should finally run the method on the empirical data, the empirical data is sometimes difficult and expensive to obtain, with unknown ground truth. On the contrary, the simulated data can be easily obtained at a low cost, and its ground truth can be under full control. These features allow the simulated data to serve as the cornerstone to benchmark new methods.

Despite the existence of more than twenty simulators for NGS technologies (Escalona *et al*., 2016), there are only three simulators created for the Nanopore sequencing, namely ReadSim (Lee *et al*., 2014), SiLiCO (Baker *et al*., 2016), and NanoSim (Yang *et al*., 2017). Although there are some differences between the three simulators (shown in Section S1), they share the same property of generating the simulated data utilizing the input nucleotide sequence and the explicit *profiles*^1^ with a statistical model. However, those simulators do not truly capture the complex nature of the Nanopore sequencing procedure, which contains multiple stages including sample preparation, current signal collection, and basecalling (shown in Fig. 1(A)). More importantly, the current signal is the essence of Nanopore sequencing, yet there is no such simulator that attempts to mimic the signal generation step.

**Fig. 1.**
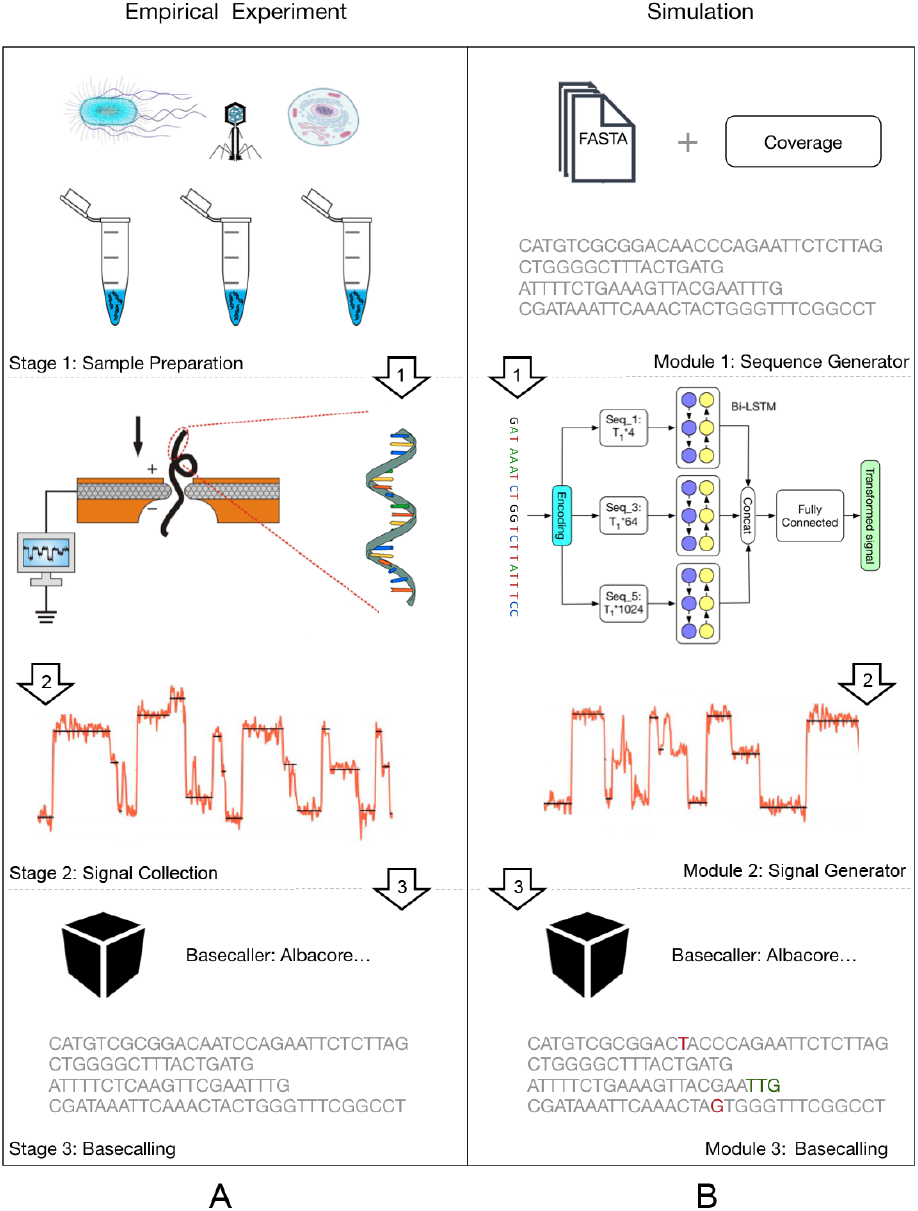
(A) The Nanopore sequencing procedure. (B) The main workflow of DeepSimulator. It simulates the entire pipeline of the empirical Nanopore sequencing experiment, producing both the simulated signals and the final simulated reads. In addition, DeepSimulator is highly modularized, which means it can be customized and updated easily to keep up with the development pace of the Nanopore sequencing technologies. Unlike the real data, the ground truth and the annotation of the simulated reads are easy to acquire. In the simulated reads on the bottom left of the figure, the red colored bases are the mismatches. The green colored bases indicate that there are indel (insertion and deletion) before them.

Instead of following the commonly adapted scenario of designing a simulator from the statistical aspect, we tackle the problem from a different angle, proposing a novel simulator that is designed more naturally for Nanopore sequencing. To run the simulator, the user just need to input a reference genome or assembled contigs, specifying the coverage or the number of reads. The sequence would first go through a preprocessing stage, which produces several shorter sequences, satisfying the input coverage requirement and the read length distribution of real Nanopore reads. Then, those sequences would pass through the signal generation module, which contains the pore model component and the signal repeating component. The pore model component is used to model the expected current signal of a given *k*-mer (k usually equals to 5 or 6 and here we use 5-mer without loss of generality), which is followed by the signal repeating component to produce the simulated current signals. These simulated signals are similar to the real signals in both strength and scale. Finally, the simulated signal would go through Albacore^2^, the ONT official basecaller, to produce the final simulated reads.

Obviously, the core component of our simulator is the pore model in the signal generation module. Currently, all the existing pore models^3^ are context-independent ones, which assign each 5-mer a fixed value for the expected current signal regardless of its locations on the nucleotide sequence. In order to further polish our simulator, we propose a novel context-dependent pore model, taking advantage of deep learning method, which has shown great potential in bioinformatics (Alipanahi *et al*., 2015; Li *et al*., 2017; Dai *et al*., 2017). Nonetheless, it is not straightforward to train the deep learning model because of the fact that the current signal is usually 8-10 times longer than the nucleotide sequence. To conquer this difficulty, we propose a novel deep learning strategy BiLSTM-extended Deep Canonical Time Warping (BDCTW), which combines bi-directional long short-term memory (Bi-LSTM) (Graves and Schmidhuber, 2005) with deep canonical time warping (DCTW) (Trigeorgis *et al*., 2016) to solve the scale difference issue.

As described above and shown in Fig. 1 (B), our DeepSimulator is “deep” in two folds. First, instead of being a simulator that only mimics the result, our simulator mimics Nanopore sequencing deeply by simulating the entire processing pipeline. Secondly, when translating the sequences into the current signals, we build a context-dependent pore model using deep learning methods. By mimicking the way Nanopore works, our simulator simulates the complete Nanopore sequencing process, producing both the simulated current signals and the final reads. Besides, employing the official basecaller, our simulator not only eliminates the procedure of learning the parameters in the profile, but also indeed deploys the actual parameters implicitly. Furthermore, by dividing the simulation procedure into several modules, our simulator offers more flexibility. For instance, the user can choose to use a different basecaller, or tune the parameters in the signal generation module to obtain the final reads with different accuracies.

In summary, the main contributions of this paper are as follows:

1. We propose the first process-based simulator, DeepSimulator, which can fully simulate the entire procedure of Nanopore sequencing, producing not only the final simulated reads but also the intermediate electrical current signals.
2. We propose a novel method to simultaneously handle the temporal alignment and the correlation analysis between the current signals and the DNA sequence that have large differences in the temporal scale. In doing so, our method is based on DCTW with Bi-LSTM as the feature mapping function for handling the sequential data.
3. We propose the first context-dependent pore model, which can accurately and specifically predict the expected current signal for each 5-mer of the DNA sequence, taking into account the sequentially contextual information.

## 2 METHODS

### 2.1 Main Workflow

The main workflow of our DeepSimulator is shown in Fig. 1. Unlike the previous simulators (Yang *et al*., 2017; Baker *et al*., 2016) that only simulate the final reads from statistical models, our simulator attempts to mimic the entire pipeline of Nanopore sequencing. There are three main stages in Nanopore sequencing. The first stage is sample preparation which would result in the nucleotide specimen used in the experiment. After obtaining the specimen, the next stage is to measure the electrical current signals of the nucleotide sequences using a Nanopore sequencing device, such as the MinION. These collected signals are usually stored in a FAST5 file. Finally, we would obtain the reads by applying a basecaller to the current signals. Correspondingly, DeepSimulator has three modules. The first module is the sequence generator. Providing the whole genome or the assembled contigs, as well as the desired coverage requirement, DeepSimulator generates relatively shorter sequences, which satisfy the coverage requirement and the length distribution of Nanopore reads. The read length distribution is described in Section 2.2. Then, those generated sequences are fed into the second module, namely the signal generation module. As the core module of DeepSimulator, it is used to generate the simulated current signals which aim to approximate the current signals produced by the MinION. There are two components within this module: the pore model component and the signal simulation component. The pore model component takes as input a nucleotide sequence and outputs the context-dependent expected current signal for each 5-mer in the sequence, which is discussed in details in Section 2.3. The signal simulation component repeats an expected signal several times at each position based on the signal repeat time distribution and then adds a random noise to produce the simulated current signals. This component is discussed in Section 2.4. The last module of DeepSimulator is the commonly used basecallers.

Notice that during the entire simulating process, we do not explicitly introduce mismatches and indels (insertions and deletions), which is usually performed in the statistical simulators (Yang *et al*., 2017; Baker *et al*., 2016) directly at the read-level. Instead, we try to mimic the current signal produced by Nanopore sequencing as similar as possible, making the basecaller introduce mismatches and indels by itself. Thus, the mismatches and indels in our method are implicitly introduced at the signal-level, which is more reasonable and closer to the realistic situation.

### 2.2 Sequence Generation

The first module of our simulator is the sequence generator. Given the user-specified reference genome or assembled contigs, as well as the desired coverage or the number of reads, the sequence generation module randomly chooses a starting position on the genome or contigs to produce the relatively short sequences, which satisfy the coverage requirement and the length distribution of the experimental Nanopore reads.

As discussed in the previous papers (Yang *et al*., 2017; Baker *et al*., 2016), the read length of Nanopore sequencing is not very straightforward to model. Many factors, such as the experimental purpose and the experimenter’s experience, would influence the read length distribution greatly. By investigating the dataset published by Nanoporetech and datasets provided by our collaborators, we find that the distribution of the read length can be categorized into three patterns by using DBSCAN (Ester *et al*., 1996) as the clustering method and histogram intersection (Swain and Ballard, 1991) as the distance metric (Fig. 2). For the first pattern shown in Fig. 2(A), we use an exponential distribution to fit it (e.g., reads for human genome). For the second pattern shown in the Fig. 2(B), we use a beta distribution to fit it (e.g., reads for *E. coli* genome). For the last pattern shown in Fig. 2(C), it is not easy to fit it using a single distribution (e.g., reads for lambda phage genome). To deal with this pattern, we use a mixture distribution with two gamma distributions to fit it. When using the simulator, the users can choose either of the three patterns. The distribution details could be referred to Section S2. Alternatively, the user can also specify the other distribution patterns for the read length.

**Fig. 2.**
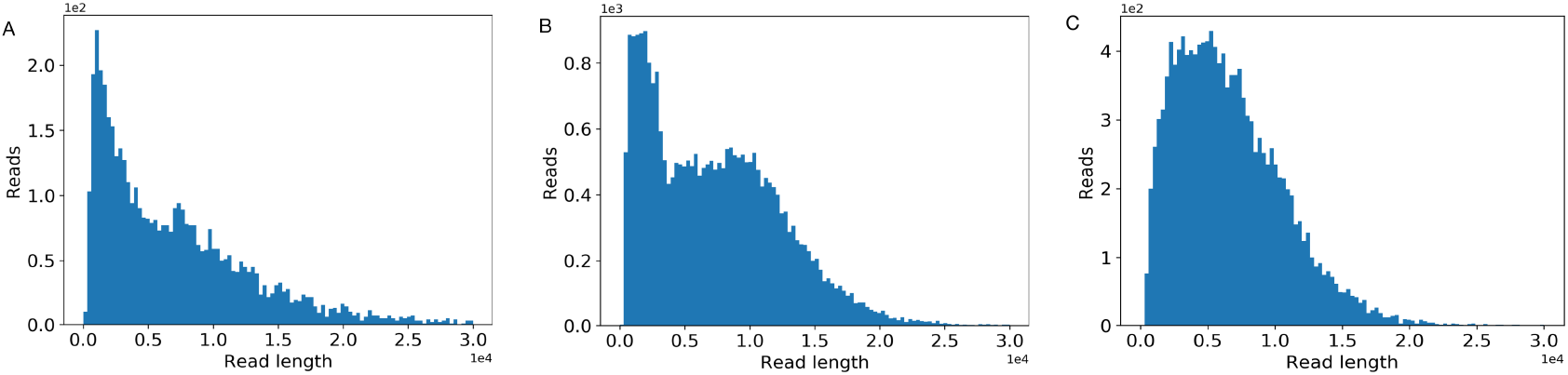
The three common read length distribution patterns in Nanopore sequencing.

### 2.3 Context-dependent Pore Model

Given a nucleotide sequence, the first step to simulate its corresponding current signals is the transformation to its expected current signals via the pore model. In this subsection, we would first formulate the problem of building the pore model, followed by the corresponding solution, BiLSTM-extended Deep Canonical Time Warping (BDCTW). We divide BDCTW into three parts: general framework of deep canonical time warping, feature representation, and neural network architecture. Finally, we introduce the productive context-dependent pore model.

#### 2.3.1 Problem formulation

A pore model is defined as the correspondence between the expected current signal and the 5-mer nucleotide sequence that is in the pore at the same time (Deamer *et al*., 2016). The pore model prediction problem is formulated as follows: given an input nucleotide sequence *X* = *x*_1_, *x*_2_,…, *x*_*T*_1__ with *T*_1_ nucleotides where *x_i_* is a 4-state nucleotide base that can take one of the four values from {A,T,C,G} for DNA or {A,U,C,G} for RNA, we need to predict the corresponding expected electrical current signals *Y* = *y*_1_, *y*_2_,…, *y*_*T*_1–4__, where *y_i_* is the predicted expected electrical current signal of a 5-mer starting from position *i* in *X* (e,g, “ACGTT”).

Here, we propose a novel method for building the pore model in consideration of the contextual information. Specifically, our method learns the context-dependent (or position-specific) pore model *Y^dep^* with length *T*_1_ – 4 for the nucleotide sequence *X* with length *T*_1_ from the raw signals (i.e., the observed current signals from a Nanopore sequencing device) *Ŷ* with length *T*_2_.

There are three challenges for learning the context-dependent pore model.

- **Scale difference.** Since the frequency of the original electrical current measurements (taken at 4000 Hz) is about 8-10 times faster than the speed at which the single-strand nucleotide sequence passes through the pore (the translocation speed is around 450 bases per second) (Stoiber and Brown, 2017), the temporal scale difference between the raw signals *Ŷ* and the nucleotide sequence *X* is large.
- **Dimensionality difference.** The feature space dimensionality is different between *X* and *Ŷ*, due to the fact that *Ŷ* is a onedimensional electrical current signal sequence whereas *X* is a nucleotide sequence with the feature dimension being at least four. This is because in order to preserve the original sequence information, one-hot encoding is commonly used (Graves, 2013) and thus four-dimension is needed to encode the four nucleotide bases.
- **Complex non-linear correlation.** The measurement of the raw signals *Ŷ* is under an extremely noisy environment because of voltage changes, noise and interactions between nanopore channels, etc (David *et al*., 2016). Thus, the relationship between *X* and *Ŷ* is very complex, having high-order or non-linear correlation.

#### 2.3.2 General framework of deep canonical time warping

The goal of deep canonical time warping (DCTW) is to discover a hierarchical or recurrent non-linear relationship between two input linearly structured data sets *X*_1_ and *X*_2_ with different lengths *L*_1_, *L*_2_ and feature dimensionality *d*_1_, *d*_2_ (i.e., *X_i_* ∈ ℝ^*d_i_* × *L_i_*^) (Trigeorgis *et al*., 2016). That is, DCTW simultaneously performs spatial transformation and temporal alignment between the two input data sequences. In our case, the two inputs are the nucleotide sequence *X* and the observed electrical current signal sequence *Ŷ*. As shown in Fig. 3, after DCTW, the transformed features from *X* and *Ŷ* are not only temporally aligned with each other, but also maximally correlated. To this end, let us consider that *Y*_i_ = *F_i_* (*X_i_*; *θ_i_*) representing the activation function of the final layer of the corresponding deep neural network (DNN) for *X_i_*, which has *d* maximally correlated units where *d* ⩽ min(*d*_1_, *d*_2_). Such an operation reduces the input data samples to the same feature dimension and then performs a maximal correlation analysis, which essentially resembles the classical canonical correlation analysis (CCA) (Akaike, 1976). Consequently, we try to optimize the following objective function,

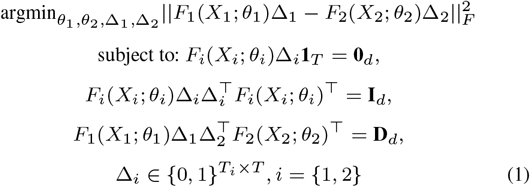

where *X*_1_ = *X* and *X*_2_ = *Ŷ*. *T*_1_, *T*_2_ and *T* are the length of *X*, *Ŷ*, and the final alignment, respectively. Δ_*i*_ are the binary selection matrices that encode the alignment paths for *X_i_*. That is, Δ_1_ and Δ_2_ remap the nucleotide sequence *X* with length *T*_1_ and raw signals *Ŷ* with length *T*_2_ to a common temporal scale *T*. **D** is a diagonal matrix. **I** is the identity matrix. And **1** (**0**) is an appropriate dimensionality vector of all 1’s (0’s).

Such an objective function can be solved via alternating optimization (Trigeorgis *et al*., 2016). Specifically, given the final layer output *F_i_* (*X_i_*; *θ_i_*), we employ dynamic time warping (DTW) (Salvador and Chan, 2007) to obtain the optimal warping matrices Δ_*i*_ which temporally align the input sequence *X_i_* and the final alignment. After obtaining the warping matrices Δ_*i*_ via DTW, we infer the maximally correlated nonlinear transformation on the temporally aligned input features *F_i_*(*X_i_*; *θ_i_*) by maximizing the following function,

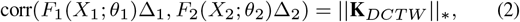

where ║·║_*_ is the nuclear norm, 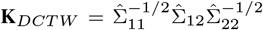 is the kernel matrix of DCTW, 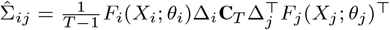 denotes the empirical covariance between the transformed data sets, where **C**_*T*_ is the centering matrix, 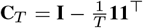.

The gradient of the objective function ║**K**_*DCTW*_║_*_ with respect to the activation layer of one neural network, such as *Y*_1_ = *F*_1_(*X*_1_; *θ*_1_), can be calculated as

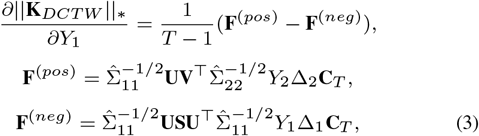

where **USV**^T^ = **K**_*DCTW*_ is the singular value decomposition (SVD) of the kernel matrix **K**_*DCTW*_. By employing this equation as the subgradient, we can optimize the parameters *θ_i_* in each neural network via back-propagation.

Since the electrical current signal of a 5-mer could be influenced by the surrounding sequences, we extend the feature function *F*_1_(*X*_1_; *θ*_1_) in the original DCTW with bi-directional long short-term memory (Bi-LSTM) (Boža *et al*., 2017) to incorporate the contextual information. Section S1 gives a brief introduction to Bi-LSTM. The DNN architecture in Fig. 3 is further elucidated in Fig. 4, which is introduced in details in Sections 2.3.3 and 2.3.4.

**Fig. 3.**
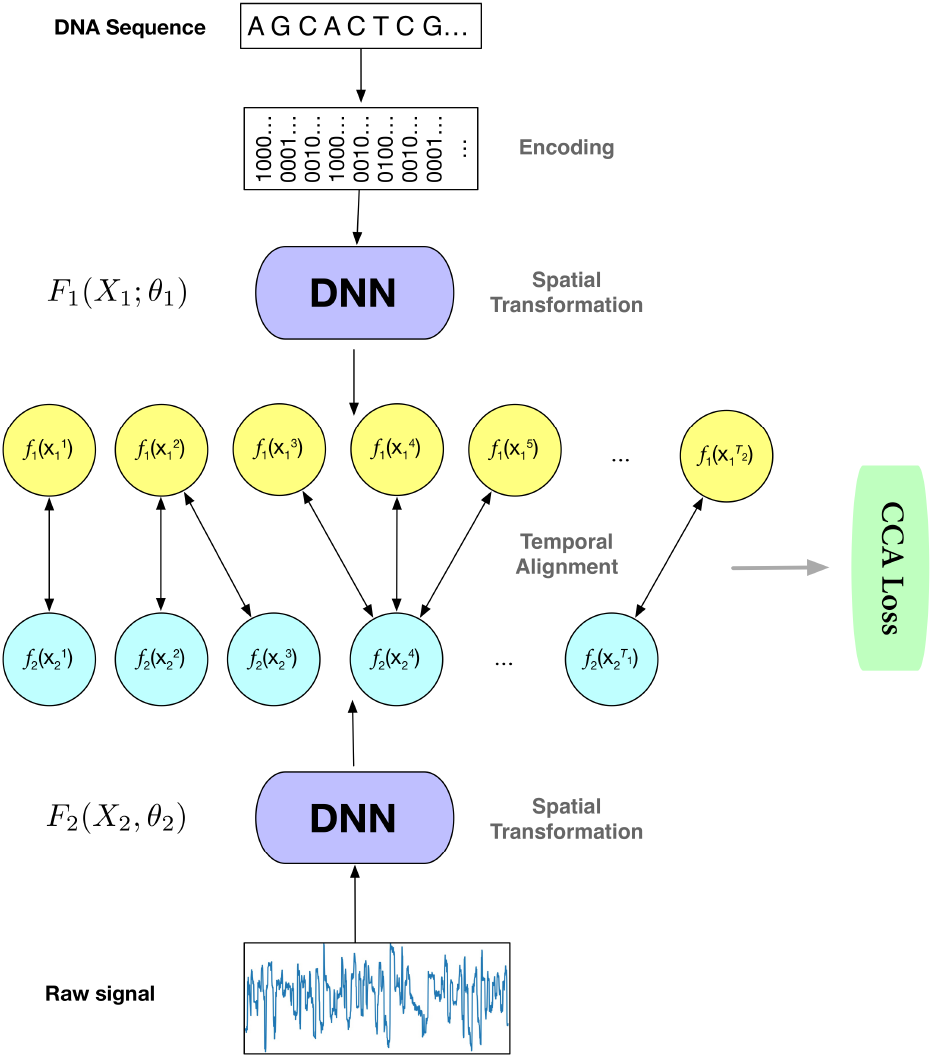
Illustration of the deep canonical time warping (DCTW) architecture with two deep neural networks (DNNs), one for the input nucleotide sequence (here we use one-hot encoding for each nucleotide and thus the feature dimension is four) and the other for the observed electrical current measurements (denoted as raw signals with feature dimension one). We train this model in an end-to-end manner, which first performs a spatial transformation that efficiently reduces the input data samples to the same feature dimension, followed by a temporal alignment that effectively maps the samples of each input sequence to a common temporal scale. The objective function of the model is to make the transformed input data samples to be maximally correlated under the canonical correlation analysis (CCA) loss.

**Fig. 4.**
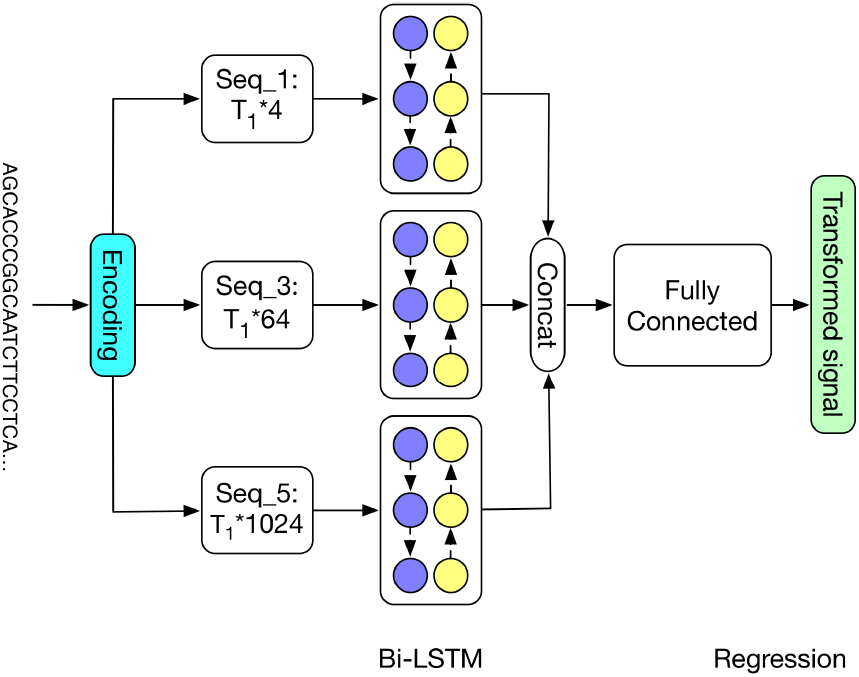
Detailed architecture of the deep neural network in deep canonical time warping for feature mapping of the input nucleotide sequence. Here we apply Bi-LSTM with three feature matrices (described in Section 2.4): Seq_*k*_ represents the feature matrix by one-hot vector encoding of *k*-mers where *k* = {1, 3, 5}, respectively. After training, this model becomes the context-dependent pore model.

#### 2.3.3 Feature representation

To preserve the original sequence information, we use one-hot encoding as the representation of the nucleotide sequence *X*. When a nucleotide sequence passes through the nanopore, each 5-mer inside the pore will cause a change in the magnitude of the electrical current. Thus, instead of just considering one nucleotide (4^1^ = 4 combinations) at position *t*, we encode the 3-mer (4^3^ = 64 combinations) and the 5-mer (4^5^ = 1024 combinations) centered at *t* as well. Specifically, we use one 1 and (4^k^ – 1) 0’s to represent each *k*-mer (*k* ∈ {1, 3, 5}). Then for each nucleotide sequence *X* with length *T*_1_, the one-hot encoding would produce three feature matrices with dimensions *T*_1_ × 4, *T*_1_ × 64, and *T*_1_ × 1024, respectively. Each row in the feature matrix represents a specific position and each column represents the appearance of a certain *k*-mer.

#### 2.3.4 Neural network architecture

To simplify our model architecture, we use an identical transformation as the feature mapping to deal with the raw signal data. That is, we set *F*_2_ (*X*_2_; *θ*_2_) = *Ŷ*. For the other feature mapping function *F*_1_ (*X*_1_; *θ*_1_) for the nucleotide sequence, we use the Bi-LSTM architecture. Specifically, as shown in Fig. 4, for each feature matrix, we use a Bi-LSTM block to obtain the hidden representation, with 50 forward LSTM cells and 50 backward LSTM cells. After concatenating the obtained hidden representation of different feature matrices, we feed it into a fully-connected layer with 200 nodes, which is followed by a regression layer. All the weights are initialized using the Xavier method. To avoid overfitting, we utilize weight decay with the coefficient as 1e^-4^. We choose Adam (Kingma and Ba, 2014) as the optimizer with the learning rate 1e^-4^. Deploying batch normalization (Ioffe and Szegedy, 2015) to accelerate the train, we set the batch size as 64 during training. The deep neural network model is implemented using Tensorflow (Abadi, 2016) and can converge within 6 hours with the help of two Pascal Titan X cards.

#### 2.3.5 Context-dependent pore model

The deep neural network in deep canonical time warping for feature mapping of the input nucleotide sequence (Fig. 4) becomes the context-dependent pore model after training. To use it, the pore model first uses one-hot vector encoding of *k*-mers, where *k*=1, 3, 5, to encode the input sequence. The encodings then go through BiLSTM layers, fully-connected layers as well as the final regression layer to generate the expected electrical signals.

### 2.4 Signal Simulation

After obtaining the expected current signals of a given nucleotide sequence, the second step of simulating its corresponding current signals is to repeat the signal at each position and add random noise. It is well-known that during sequencing, the raw signal acquisition speed is much faster than the DNA or RNA moving speed, causing a certain 5-mer being measured multiple times. Thus, to convert the expected signals produced by the pore model to the current signals which can be put into a basecaller, we need to repeat a certain signal several times. Similar to the read length, we manage to model the repeat time using a mixture alpha distribution. When running the simulator, the repeat time would be drawn from the distribution for each position on the expected signal, generating the simulated current signal by repeating that position for a certain number of times. The details of the distribution and the parameters could be referred to Section S2. It should also be noted that the raw signals are extremely noisy due to the complicated sequencing environment, including voltage changes, noise and interactions between channels (David *et al*., 2016). Therefore, we add Gaussian noise with the user-defined variance parameter to each position of the simulated signals.

The main difficulty of this step is to get the statistics of the repeat time, as shown in Fig. 5. Currently, it is almost impossible to get the precise repeat time of a certain 5-mer, but it is possible to obtain the approximate repeat time statistics. Here we show the four basic steps for obtaining the statistics. (i) Taking as input the reference genome, raw signals produced by the MinION, and the basecalled reads from Albacore, we first map the reads on to the reference genome by Minimap (Li, 2016), which would mark out the ground truth (at least approximate) sequence that corresponds to the raw signal. (ii) With the ground truth sequence, we can get the expected signal of each 5-mer in the sequence using the context-independent pore model. (iii) We then apply dynamic time warping (DTW) (Salvador and Chan, 2007) to map the raw signal and the expected signal, which is based on the fact that those two signals should have the similar shape. (iv) Based on the mapping, we can find out the repeat time from the raw signal positions that correspond to each expected signal position. Performing the above procedure on a large dataset, we can get a stable statistic of the repeat time. We then fit the distribution as a mixture model (Section S3).

**Fig. 5.**
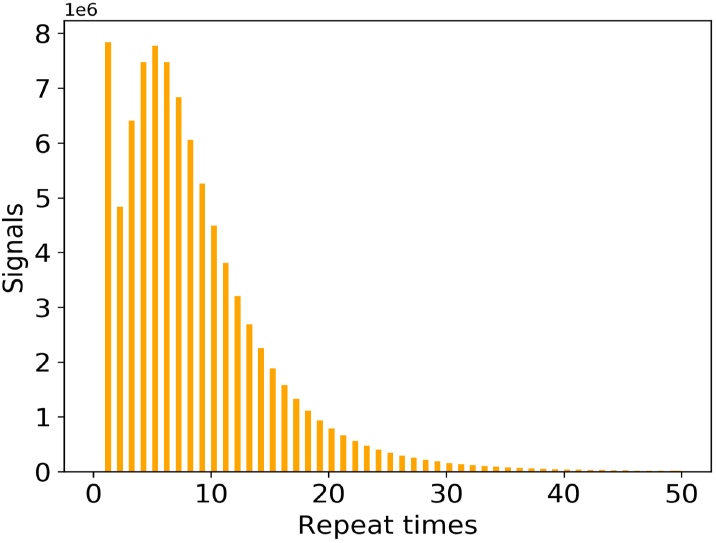
The distribution of the signal repeat times of 5-mer nucleotides.

### 2.5 Datasets

Four Nanopore sequencing datasets from different species are used in this paper: ranging from the in-house datasets lambda phage, *E.coli* K-12 sub-strain MG1655, *Pandoraea pnomenusa* strain 6399, to the public available human data. The three in-house datasets were prepared and sequenced by Prof. Lachlan Coin’s lab at University of Queensland. In particular, all the samples were sequenced on the MinION device with 1D ligation kits on R9.4 flow cells (SQK-LSK108 protocol). The publicly available human dataset is the human chromosome 21 from the Nanopore WGS Consortium (Jain *et al*., 2017b). The samples in this dataset were sequenced from the NA12878 human genome reference on the Oxford Nanopore MinION using 1D ligation kits (450 bp/s) with R9.4 flow cells. The Nanopore raw signal datasets in the FAST5 format were downloaded from nanopore-wgs-consortium^4^. The reference genomes of the four datasets were downloaded from NCBI^5^.

The context-dependent pore model of the second module in DeepSimulator was trained on the *Pandoraea pnomenusa* dataset. To construct the dataset used in Section 3.2, which is used to check the performance of the pore models, we randomly sampled 700 reads from each of remaining three species to form a dataset containing 2100 reads.

In addition to the four species for which we have both the reference genome and the empirical experimental data, we also include another extremely small genome, mitochondria, for which we only have the reference genome^6^. We used the *E.coli* K-12 genome, the lambda phage genome, and the mitochondrial genome to perform the assembly experiments in Section 3.4. Finally, the mitochondrial genome and lambda phage genome were used for the SNP calling experiments in Section 3.5.

## 3 RESULTS

We comprehensively evaluated each of the three modules in DeepSimulator. In summary, the results in this section show that (i) the length distribution of the simulated reads satisfies the empirical read length distribution; (ii) the signals generated by our context-dependent pore model are more similar to the experimental signals than the signals generated by the official context-independent pore model; and (iii) the final reads generated by DeepSimulator with the default parameter have almost the same profile as the experimental data. We finally show that DeepSimulator can benefit the development of tools or methods in *de novo* assembly and low coverage SNP detection.

### 3.1 Read Length Distribution

As mentioned in Section 2.2, for an input genome sequence, DeepSimulator generates reads whose length distribution satisfies the empirical length distribution. We provide three predefined distributions, beta distribution, exponential distribution, and the mixed gamma distribution, which cover the three main patterns of the Nanopore read length distribution (Fig. 2). The parameters of these distributions are given in Section S2. In general, the mixed gamma distribution is often the most suitable length distribution. As a result, we set it as the default length distribution pattern. In addition to that, considering the property of different sequencing tasks, some biological experiments may be designed on purpose so that the read length distribution would satisfy a predefined distribution. In order to simulate this case, we also provide the interface for the user-defined read length distributions. The distributions of the length of the simulated reals by DeepSimulator on human, *E.coli* K-12 sub-strain MG1655, and lambda phage are very similar to that of the experimental reads (Section S4).

### 3.2 Simulated Signals

To check the signal-level similarity between the simulated signals generated by DeepSimulator and the experimental ones produced by the MinION (i.e., the raw signals), we employed dynamic time warping (DTW) (Salvador and Chan, 2007) (see Section S5 for details) which is the standard way of checking the difference between two signal sequences on the randomly selected 2100 reads from lambda phage, *E.coli* K-12 substrain MG1655, and human (Section 2.5). The average deviation between them is 0.175. We also performed the same analysis using the official content-independent pore model followed by the same signal repeat component used in DeepSimulator to obtain the context-independent simulated signals. Using the same set of reads, the average deviation of the context-independent signals to the raw ones is 0.185, which is about 5.7% higher than that of DeepSimulator. Furthermore, we performed another experiment on the reads generated by NanoSim (Yang *et al*., 2017) to derive the simulated signals by the context-independent pore model. The average deviation of the NanoSim signals to the raw ones is 0.210, which is 20% higher than that of DeepSimulator. Fig. 6 shows the comparison of the deviation scores of the DeepSimulator signals and that of the context independent signals as well as that of the NanoSim signals for the 2100 reads. Notice that DeepSimulator was trained solely on *Pandoraea pnomenusa* and tested on the three other species, which demonstrates the generality of our model.

**Fig. 6.**
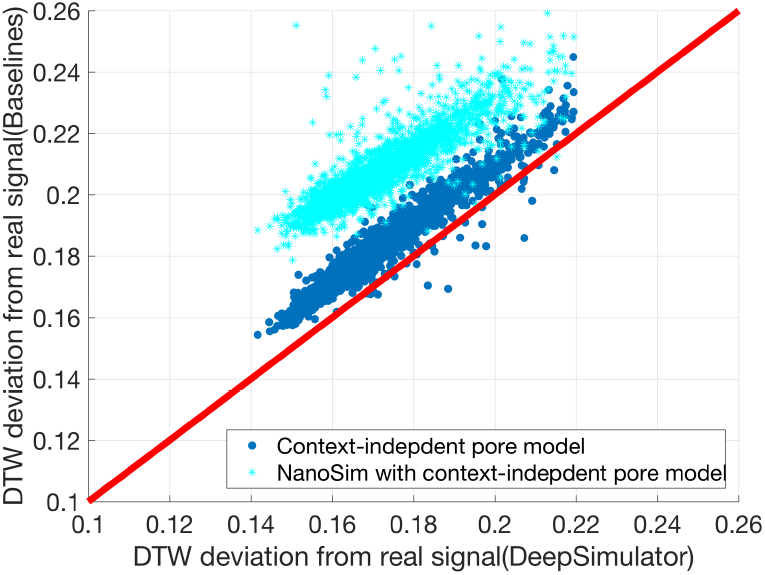
Comparison of the context-dependent pore model component of DeepSimulator with the context-independent pore model on the signal-level. Each point represents an input read. The x-axis represents the DTW deviation of the DeepSimulator signals of the input read from the real raw signals. The y-axis represents the DTW deviation of the signals generated from context-independent pore model from the real raw signals (context-independent pore model with our signal repeat component in blue, and context-independent pore model with NanoSim in cyan). The red line is the diagonal line. Any point above the red line means our simulation is better, whereas any point below means the existing method is better.

### 3.3 Simulated Reads

For the read-level outputs, we provided a parameter interface in DeepSimulator, which can be adjusted continuously so that the user could control the final read basecalling accuracy as well as the indel ratio. Internally, the parameters change the noise and the signal repeat time distribution, which are the two factors that affect the read profile greatly. To check the read profile of the simulated reads, for a given input ground truth sequence, we ran DeepSimulator to obtain the simulated read. Performing BLAST (Altschul *et al*., 1997) between the simulated read and the input ground truth read, we can calculate the profiles such as the accuracy, mismatch number, and gap numbers. According to our experiment, the output reads of DeepSimulator can have a basecalling accuracy ranging from 83% to 97%. Table 1 shows the profile of the real reads and the profiles of DeepSimulator reads using four typical parameter settings. In addition, we also checked the profile of the reads generated from the official context-independent pore model, whose output is extended using the noise-free repeat time distribution and further basecalled using Albacore, which is shown in the fourth column of Table 1.

**Table 1.** The profiles of different types of reads, which are basecalled using Albacore. DS represents the reads generated from DeepSimulator. Here we show the profiles of four typical settings (the parameter can be adjusted continuously, not only just four choices) of DeepSimulator, noise free, high accuracy, middle accuracy (aimed at simulating the emperical data profile), and low accuracy. OPM (official pore model) shows the read profile generated by the official context-independent pore model, whose output is extended using the noise-free repeat time distribution and further basecalled using Albacore, given an input ground truth sequence. The parameter manual of DeepSimulator can be referred to Section S6.

| Criteria | Real data | OPM | DS (noise free) | DS (high acc) | DS (med acc) | DS (low acc) |
| --- | --- | --- | --- | --- | --- | --- |
| Accuracy | 88.49 | 95.99 | 97.01 | 92.96 | 88.78 | 83.45 |
| Mismatch | 2.88 | 1.24 | 0.94 | 1.87 | 2.74 | 4.36 |
| Gap open | 5.38 | 2.21 | 1.69 | 3.63 | 5.28 | 7.08 |
| Gap total | 8.62 | 2.77 | 2.04 | 5.17 | 8.48 | 12.19 |

### 3.4 *De novo* Assembly

Because of long reads, Nanopore sequencing has higher potential in genome assembly than the other short-reads sequencing technologies. Thus, one of the main applications for Nanopore sequencing is *de novo* assembly. We used two widely recognized *de novo* assembly pipelines, Canu (Koren *et al*., 2017) and Miniasm (Li, 2016) with Racon (Vaser *et al*., 2017), to perform such task on two different sets of simulated reads generated by DeepSimulator from the *E.coli* K-12 genome and the lambda phage genome, respectively. Both experiments succeed in assembling the simulated reads into one contig. The comparison between the assemblies and the reference genome is plotted using MUMmer (Delcher *et al*., 1999), as shown in Fig. 7(A, B). As a comparison, we also show the assembly results of *E.coli* K-12 and lambda phage using the empirical data. Fig. 7(C, D) illustrate that the results of the empirical data show similar patterns as the results of the simulated data. In addition to the relatively large genome, *E.coli* K-12, which is 4.6 Mbp, and a small genome, lambda phage, which is 48 Kbp, we also performed another experiment on an extremely small genome, the mitochondrial genome (16 Kbp). Miniasm with Racon also succeeded in assembling the simulated reads into one contig (Section S7).

**Fig. 7.**
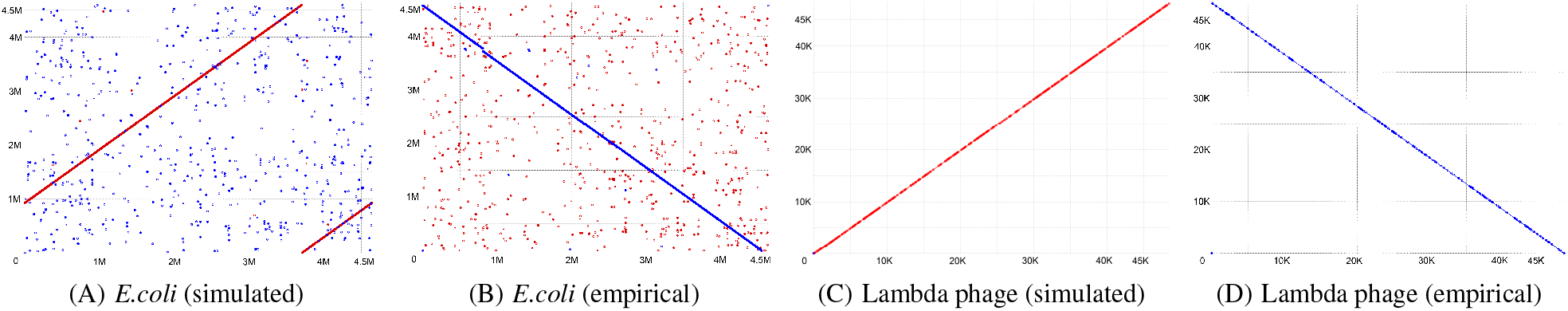
Mummer plots comparing the reference genome on the x-axis with the assembled genome on y-axis. (A) The assembly result of *E.coli* K-12 genome by Canu, using simulated reads from DeepSimulator. (B) The assembly result of *E.coli* K-12 genome by Canu, using the experimental MinION sequence data (i.e., empirical data). (C) The assembly result of lambda phage genome by Miniasm with Racon, using simulated reads from DeepSimulator. (D) The assembly result of lambda phage genome by Miniasm with Racon, using the empirical data.

### 3.5 Low Coverage SNP Detection

Single nucleotide polymorphisms (SNPs) are found to be involved in the etiology of many human diseases. For example, hundreds of SNPs in the mitochondrial DNA (mtDNA) have been linked to aging-related diseases (Stewart and Chinnery, 2015; Ocampo *et al*., 2016). Despite the importance of the complete haplotyping of the mitochondrial genome, the current methods, which are designed for detecting mitochondrial mutations from a population of cells, would perform massively parallel sequencing of short DNA fragments, having difficulty in performing the complete haplotyping. On the other hand, the Nanopore sequencing, which has the potential of performing the long-read single-molecular sequencing of mtDNA, may overcome the hurdle. Under this circumstance, mimicking the ideal single molecular Nanopore sequencing scenarios, we conducted experiments on the success rate of SNPs detection with respect to sequencing coverage, using the simulated reads from DeepSimulator.

Considering the basecalling accuracy of the Nanopore sequencing, although the current basecalling accuracy is not high enough (around 86% to 88%), theoretically, we can consider those errors as random errors instead of systematic errors, and the consensus analysis could help us get rid of such random noise and detect the systematic variants which are caused by SNPs.

The results are shown in Fig. 8. On the simulated data of mitochondrial genome, we could detect SNPs when the coverage is above 6X using the standard pipeline of samtools (Li *et al*., 2009) and bcftools (Li, 2011) (Fig. 8(A)), which is consistent with the conclusion in (Zeng *et al*., 2013). As the number of the implanted SNPs increases, the coverage should increase to ensure all the SNPs to be successfully called. Fig. 8(B) shows the same analysis on the lambda phage genome, which shares the similar pattern as the mitochondrial experiment. In summary, the detection of the SNPs would become more difficult as the number of SNPs increases. Our experiments demonstrate that in general, 6× coverage would be enough to detect a small number of SNPs.

**Fig. 8.**
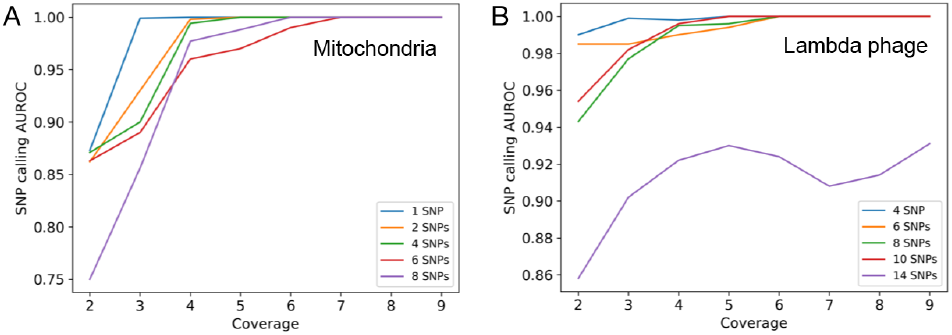
(A) The relationship between the SNP detection performance and the coverage as well as the number of introduced SNPs on the simulated reads from the mitochondrial genome. (B) The relationship between the SNP detection performance and the coverage as well as the number of introduced SNPs on the simulated reads from the lambda phage genome.

## 4 DISCUSSION & CONCLUSION

In this paper, we proposed DeepSimulator, the first Nanopore simulator that aims at mimicking the entire procedure of Nanopore sequencing. Unlike the previous simulators which only simulate the reads from the statistical patterns of the real data, DeepSimulator simulates both the raw electrical current signals and nucleotide reads.

There are three advantages of DeepSimulator. First of all, our pipeline is highly modularized, which is easier to be customized by users. For example, the users can use another basecaller, to replace Albacore, to obtain the reads with the profile of that basecaller. Secondly, because of the modularization, compared with other simulators, it is more likely for our simulator to keep up with the rapid development of the Nanopore sequencing technology. If one step of the Nanopore sequencing pipeline is updated, we can also update the corresponding module without changing the entire pipeline completely. Thirdly, in addition to the final simulated reads, we are also able to obtain the simulated electrical current signals, which are very useful for the development of basecallers and for the benchmarking of signal-level read mappers.

There are two potential applications of DeepSimulator. First, DeepSimulator can generate benchmark datasets to evaluate the newly developed methods for Nanopore sequencing data analysis. Unlike the empirical datasets whose ground truth is difficult to obtain, DeepSimulator can be fully controlled, which makes it a practical complement to the empirical data. Second, as shown in the SNP detection experiments, it can act as a guidance to the empirical experiment by simulating the ideal case.

## ACKNOWLEDGMENTS

We thank Minh Duc Cao, Lachlan J.M. Coin, Louise Roddam, and Tania Duarte for providing the nanopore sequencing data for the lambda phage, *E. coli*, and *Pandoraea pnomenusa* samples. This work was supported by the King Abdullah University of Science and Technology (KAUST) Office of Sponsored Research (OSR) under Awards No. URF/1/1976-04, URF/1/2602-01, and URF/1/3007-01.

## Footnotes

1 Here the profiles refer to a set of parameters, such as insertion and deletion rates, substitution rates, read lengths, error rates and quality scores. For instance, ReadSim uses the fixed profile; SiLiCO uses the user provided profile; and NanoSim uses the user provided empirical data to learn the profile which would be used in the simulation stage.

2 https://community.nanoporetech.com/protocols/albacore-offline-basecalli/v/abec_2003_v1_revad_29nov2016/linux

3 https://github.com/nanoporetech/kmer_models

4 http://s3.amazonaws.com/nanopore-human-wgs/rel3-fast5-chr21.part03.tar

5 https://www.ncbi.nlm.nih.gov/nuccore/J02459, https://www.ncbi.nlm.nih.gov/nuccore/U00096, https://www.ncbi.nlm.nih.gov/nuccore/JTCR01000000, https://www.ncbi.nlm.nih.gov/nuccore/NC000021

6 https://www.ncbi.nlm.nih.gov/nuccore/AY172335

